# Local rather than global H3K27me3 dynamics associates with differential gene expression in *Verticillium dahliae*

**DOI:** 10.1101/2021.05.04.442700

**Authors:** H. Martin Kramer, Michael F. Seidl, Bart P.H.J. Thomma, David E. Cook

## Abstract

Differential growth conditions typically trigger global transcriptional responses in filamentous fungi. Such fungal responses to environmental cues involve epigenetic regulation, including chemical histone modifications. It has been proposed that conditionally expressed genes, such as those that encode secondary metabolites but also effectors in pathogenic species, are often associated with a specific histone modification, lysine27 methylation of H3 (H3K27me3). However, thus far no analyses on the global H3K27me3 profiles have been reported under differential growth conditions in order to assess if H3K27me3 dynamics governs differential transcriptional. Using ChIP- and RNA-sequencing data from the plant pathogenic fungus *Verticillium dahliae* grown in three *in vitro* cultivation media, we now show that a substantial number of the identified H3K27me3 domains globally display stable profiles among these growth conditions. However, we do observe local quantitative differences in H3K27me3 ChIP-seq signal that associate with a subset of differentially transcribed genes between media. Comparing the *in vitro* results to expression during plant infection suggests that *in planta*-induced genes may require chromatin remodelling to achieve expression. Overall, our results demonstrate that some loci display H3K27me3 dynamics associated with concomitant transcriptional variation, but many differentially expressed genes are associated with stable H3K27me3 domains. Thus, we conclude that while H3K27me3 is required for transcriptional repression, it does not appear that transcriptional activation requires global erasure of H3K27me3. We propose that the H3K27me3 domains that do not undergo dynamic methylation may contribute to transcription through other mechanisms or may serve additional genomic regulatory functions.

## INTRODUCTION

The fungal kingdom comprises a plethora of species occupying an enormously diverse range of ecological niches (1). As environments are typically dynamic, including the effects of daily and yearly cycles, fungi continuously need to respond and adapt to survive (2, 3). To this end, fungi have evolved various mechanisms to monitor their environment and to transcriptionally respond to environmental cues (4). For instance, the yeast *Saccharomyces cerevisiae* senses cold stress by increased membrane rigidity, which leads to transcription of genes that, among others, encode cell damage-preventing proteins (5). Furthermore, *S. cerevisiae* senses the quantitative availability of carbon and nitrogen sources in the environment to determine which developmental program maximises the potential for survival (6). In animal systems, epigenetic mechanisms (i.e. those affecting genetic output without changing the genetic sequence) are implicated in transcriptional responses to changing environments (7–9). Such epigenetic mechanisms have similarly been proposed to contribute to environmental response in filamentous fungi. For instance, the saprotrophic fungus *Neurospora crassa* phenotypically reacts to environmental stimuli, such as changes in temperature and pH, yet *N. crassa* mutants impaired in transcription-associated epigenetic mechanisms display reduced growth in response to these stimuli (10). Similarly, the nectar-feeding yeast *Metschnikowia reukaufii* and the ubiquitous fungus *Aureobasidium pullulans* fail to properly respond to changing carbon sources when DNA methylation or histone acetylation are inhibited (11, 12). These results suggest that epigenetic mechanisms are important for transcriptional responses to changing environments in diverse fungi, but many questions remain regarding the precise mechanisms and function of epigenetic dynamics in fungi.

Epigenetic mechanisms, such as direct modifications of DNA and histone proteins or physical changes to the chromatin architecture, can influence transcription by regulating DNA accessibility (13). Chromatin that is accessible and potentially active is termed euchromatin, while heterochromatin is condensed and often transcriptionally silent (14). However, heterochromatic regions are not always repressed. Heterochromatin is subcategorized into constitutive heterochromatin that remains condensed throughout the cell cycle, and facultative heterochromatin that can decondense to allow transcription in response to developmental changes or to environmental stimuli (15–17). In fungi, constitutive heterochromatin is often associated with repeat-rich genome regions and is typically characterized by tri-methylation of lysine 9 on histone H3 (H3K9me3), while facultative heterochromatin is characterized by tri-methylation of lysine 27 on histone H3 (H3K27me3) (18–21). Empirically, both H3K9me3 and H3K27me3 have been implicated in transcriptional regulation in various fungi (22, 23, 32, 24–31). The majority of these studies rely on genetic perturbation of the enzymes that deposit methylation at H3K9 and H3K27, and the results consistently show that depletion of methylation at these lysine residues mainly results in transcriptional induction. However, as global depletion of a histone modification can result in pleiotropic effects, such as improper localization of other histone modifications or altered development (33, 34), it is difficult to infer transcriptional control mechanisms used for natural gene regulation from these genetic perturbation experiments. Therefore, additional research is needed to directly test the hypothesis that heterochromatin-associated histone modifications directly regulate transcription either through their dynamics, or their action to form or recruit transcriptional complexes (35–37).

The filamentous fungus *Verticillium dahliae* is a soil-borne broad host-range plant pathogen that infects plants through the roots to invade the xylem vessels and cause vascular wilt disease (38). Genomic and transcriptomic studies have revealed that the *V. dahliae* genome harbours lineage-specific (LS) regions that are variable between strains and enriched for genes that are *in planta* induced (39, 40). These LS regions are generally considered genomic hotspots for evolutionary adaptation to plant hosts (39–44). Recently we explored the epigenome of *V. dahliae* and distinguished LS regions from the core genome based on particular chromatin signatures, including elevated levels of H3K27me3 accompanied with accessible DNA and active transcription (44). Using a machine learning approach and supported by orthogonal analyses, we identified nearly twice as much LS DNA as previously considered, collectively referred to as adaptive genomic regions (44). Given the elevated levels of H3K27me3 at adaptive genomic regions in *V. dahliae*, and previous reports that removal of H3K27me3 results in transcriptional induction (22–24), we now tested if H3K27me3 dynamics are required for transcriptional activation of genes under different growth conditions. Ideally, the involvement of H3K27me3 dynamics in differential gene expression of *V. dahliae* would be studied between *in vitro* and *in planta* growth, as adaptive genomic regions are enriched for *in planta* induced genes (40, 44). However, *V. dahliae* displays a low pathogen-to-plant biomass during infection (45), which impedes technical procedures to determine histone modification levels over the genome (46). Nevertheless, as H3K27me3 is generally reported to regulate transcription in response to environmental stimuli (15, 17), we hypothesize that it may be involved in transcriptional regulation in *V. dahliae* during differential growth conditions *ex planta* as well. Here, we analyse RNA-seq, ChIP-seq and ATAC-seq data of *V. dahliae* cultured in various axenic growth media to understand if transcriptional dynamics require concomitant changes H3K27me3 modification status.

## RESULTS

### Chromatin features correlate with gene expression levels

To determine how general features of chromatin, such as histone modifications and DNA accessibility, impact transcriptional activity in *V. dahliae*, we mapped the occurrence of heterochromatic histone marks H3K9me3 and H3K27me3, euchromatic histone marks H3K4me2 and H3K27ac, and chromatin accessibility determined by assay for transposase accessible chromatin followed by sequencing (ATAC-seq) (47) (Fig. 1A, Fig. S1). Grouping the *V. dahliae* genes into five expression quintiles, from quintile 1 containing the highest 20% expressed genes to quintile 5 with the 20% lowest expressed genes, we are able to integrate histone modification profiles and transcriptional activity. Expressed genes (quintiles 1-4) displayed low H3K4me2 and H3K27ac coverage upstream of the transcription start site (TSS), followed by a steep increase of coverage over the start of the gene that decreases over the gene body and increases again at the transcription end site (TES) (Fig. 1B). The strength of the association corresponds to the level of transcription, also within quintiles (Fig. S2). The low H3K4me2 and H3K27ac coverage directly upstream of the TSS coincides with increased chromatin accessibility, where higher expressed quintiles have more open DNA (Fig. 1B, Fig. S2). This chromatin profile upstream of the TSS suggests occurrence of a nucleosome depleted region. There is little evidence of H3K9me3 over gene bodies and cis-regulatory regions (Fig. 1B, Fig. S2), which corroborates that H3K9me3 marks TEs in constitutive heterochromatic regions such as the centromeres (18, 44, 48, 49). During cultivation in PDB, H3K27me3 is mainly present on genes that are lowly or not expressed (quintile 4 and 5) (Fig. 1B, Fig. S2). These results, and association between chromatin features and transcriptional activity, are consistent with reports for other fungi (24, 27, 32).

**FIG 1.**
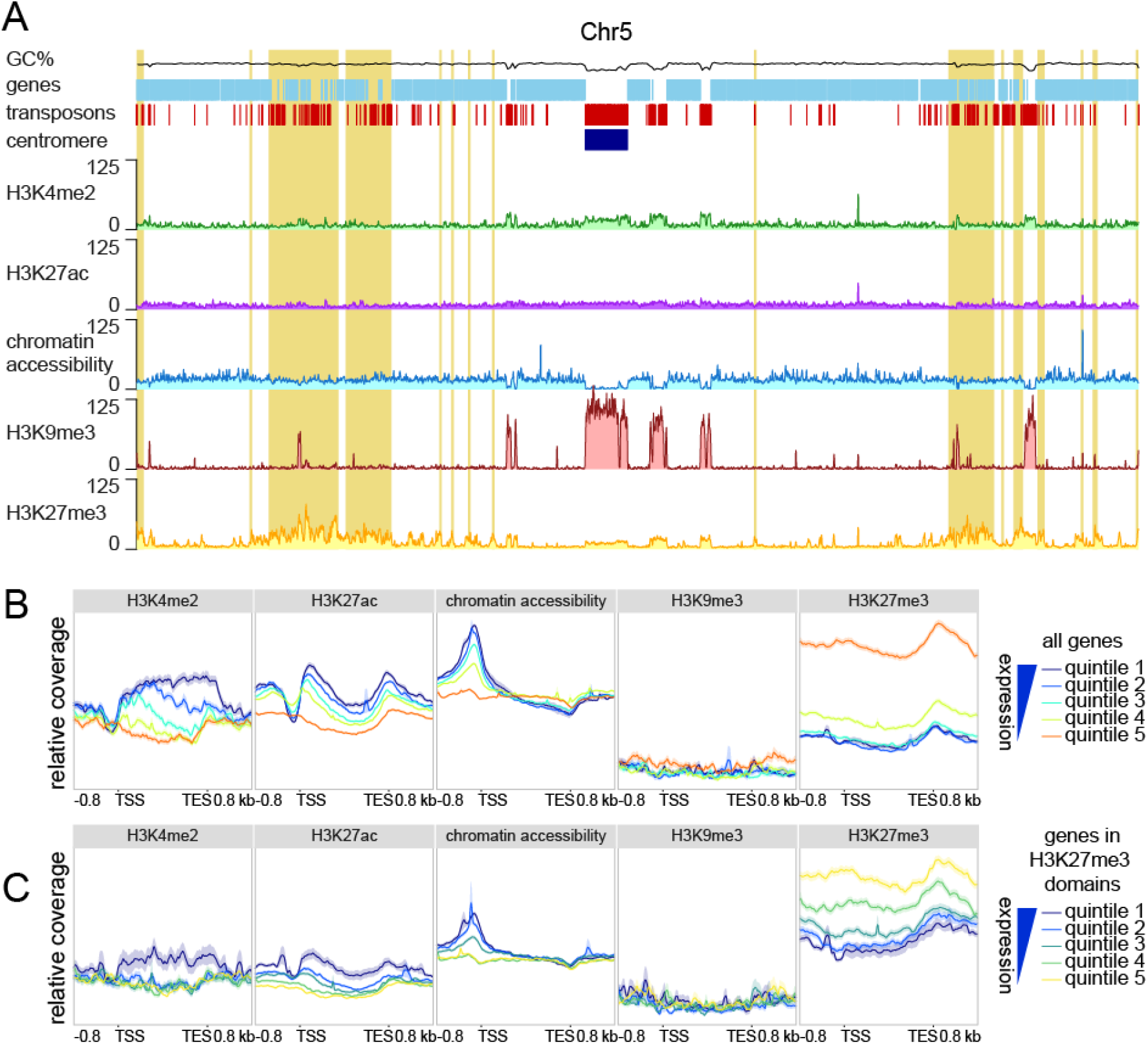
Lowly and non-expressed genes associate with H3K27me3. A) Whole genome distribution of the euchromatin-associated histone modification H3K4me2 and H3K27ac, the constitutive heterochromatin-associated histone modification H3K9me3, the facultative heterochromatin-associated histone modification H3K27me3 and chromatin accessibility as determined by ATAC-seq, based on chromosome 5 as an example. GC percentage is indicated in black, genes are indicated in light blue, transposons are indicated in red, the centromere is indicated in dark blue and adaptive genomic regions are indicated in yellow. B,C) Relative coverage of chromatin accessibility and the histone marks H3K4me2, H3K9me3, H3K27me3 over gene bodies (between transcription start site (TSS) and transcription end site (TES)) ±800 bp of flanking sequence, grouped into quintiles based on gene expression levels upon cultivation for six days potato dextrose broth (PDB), for (B) all genes, and for (C) all genes located in an H3K27me3 domain.

To further analyse the association between H3K27me3 and gene expression, we identified 3,186 genes covered by H3K27me3 peaks (see methods) from triplicate grown *V. dahliae* in PDB. These 3,186 genes were separated into five expression quintiles as previously described. Interestingly, we found that genes in H3K27me3 domains with higher expression values (quintile 1 and 2) had higher H3K4me2 over the gene body, more accessible promoters, and lower H3K27me3 values (Fig. 1C). This suggests that genes in H3K27me3 domains are not uniformly repressed or heterochromatic, and there appears to be a quantitative, rather than qualitative, association between H3K27me3 association and gene activity. Genes with a lower association to H3K27me3 may represent loci that are not in a stable state under the tested conditions, where some cells have a more euchromatic profile and other have a more heterochromatic profile. While we cannot infer these details from the current data, it is clear that genes with lower expression in PDB are generally marked with H3K27me3, have less H3K4me3, and have less accessible DNA in the region of transcription initiation (i.e. their promoter).

### Genetic perturbation of H3K27me3 induces transcription of many genes that are differentially expressed *in vitro* and *in planta*

To further characterize the influence of H3K27me3 on gene expression in *V. dahliae*, we deleted the histone methyltransferase component of the Polycomb Repressive Complex 2 (PRC2), termed **Set7** (*Δ*Set7**), leading to loss of H3K27me3 (Fig. 2A, Fig. S3, S4). We do note that some background signal is present for the H3K27me3-ChIP-seq conducted in Δ*Set7*, but the signal is relatively uniform across the genome and does not correspond to the regions of H3K27me3 found in wild-type (Fig. 2A). As H3K27me3 is generally associated with facultative heterochromatin, we anticipated that the loss of H3K27me3 would mainly lead to induction of genes that were located in H3K27me3 domains in wild-type *V. dahliae*. Out of the 839 genes that are induced in Δ*Set7* (log_2_-fold change >2, p<0.05), 625 (74.5%) locate in H3K27me3 domains, which is significantly higher than expected given that only 27.9% of genes locate in H3K27me3 domains (Fisher’s exact test, p<0.00001). In contrast, we find that 211 (27.6%) of 765 repressed genes in Δ*Set7* are in H3K27me3 domains (no association, Fisher’s exact test, p=0.94) (Fig. 2B,C). Additionally, when comparing log_2_-fold changes in expression between Δ*Set7* and the wild type strain, we observed that genes located in a H3K27me3 domain in wild type are more significantly induced in Δ*Set7* than genes not locating in H3K27me3 domains (Two-sample Student’s T-test, p<0.001) (Fig. S5). These findings support the role of H3K27me3 in transcriptional repression, and show that loss of H3K27me3 can lead to de-repression during growth *in vitro*.

**FIG 2.**
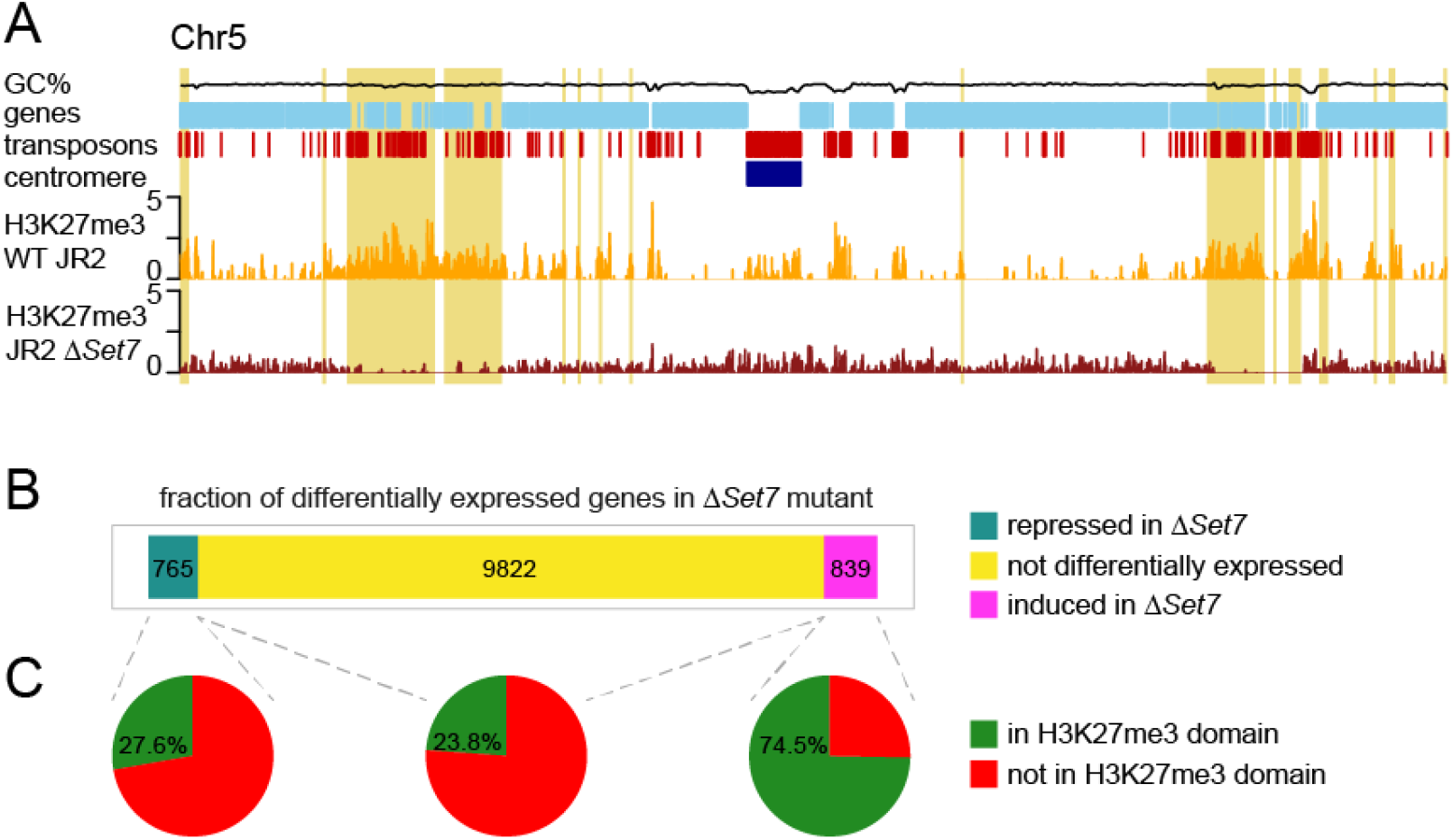
Genetic perturbation of H3K27me3 results in induction of genes that are transcriptionally regulated in different growth conditions. A) H3K27me3 ChIP sequencing on triplicate JR2 WT (yellow) and duplicate Δ*Set7* (red) samples with coverage over chromosome 5 as an example. Adaptive genomic regions are indicated in yellow. B) Fractions of induced (log2 fold change >1, p<0.05) and repressed (log2 fold change <−1, p<0.05) genes and genes that are not differentially expressed between wild-type and Δ*Set7* that (C) locate in H3K27me3 domains.

### H3K27me3 domains are globally stable between *in vitro* growth conditions

Given that H3K27me3 domains contribute to transcriptional repression, a key question concerns the status of H3K27me3 under growth conditions where the underlying genes are transcribed. One hypothesis is that H3K27me3 is removed or lost under growth conditions that activate gene expression, which would be noticeable as a change in H3K27me3 ChIP-seq domains between *V. dahliae* growth conditions that lead to differential expression. Here, the observed changes in H3K27me3 domains should be associated with transcriptional differences of the underlying genes. An alternative hypothesis is that H3K27me3 domain status does not change in accordance with transcriptional activity, and the repressive effects of H3K27me3 are released through alternative means. To test this hypothesis, we performed triplicates of H3K27me3 ChIP-seq on *V. dahliae* cultivated for six days in MS and CZA, in addition to the previously generated ChIP-seq data in PDB. Based on correlation between replicates (Fig. S6), we decided to continue with 3 replicates of H3K27me3 ChIP data in PDB and 2 replicates each of H3K27me3 ChIP data in CZA and MS. Control ChIP-input samples were used to normalize H3K27me3 datasets and identify H3K27me3-domains in each of the three growth media. We identified a total of 2,654 genes that were always present in H3K27me3 domains, regardless of the *in vitro* growth media (Fig. 3A). Interestingly, the 2,654 genes present in stable H3K27me3 domains display significantly stronger differential expression between all pair-wise media comparisons when compared with non-marked genes in all three growth conditions (Fig. 3B). This suggests that differential gene expression can occur without changes in global H3K27me3 coverage. We further checked whether genes that are differentially expressed between *in vitro* growth media are associated with H3K27me3 during cultivation in the growth medium with low expression levels (Fig. S7). Interestingly, we observed that for all pair-wise comparison between growth media, genes with higher log2 fold-changes in expression are more likely to locate in an H3K27me3 domain in the non-transcriptionally permissive growth medium (Fig. S7), again suggesting that H3K27me3 is involved in regulation of differential gene expression. To assess if changes in H3K27me3 domains between the media were associated with changes in transcription, we compared differences in domains and differential gene expression between media. We first analysed differential expression for genes present in H3K27me3 domains in two media but not in an H3K27me3 domain in the third medium. For example, we identified 366 genes that were in a H3K27me3 domain in PDB and MS but not during CZA growth (Fig. 3A), and found that these genes are not differential expressed between PDB and MS (one-sample t-test, p 0.99), but they are higher expressed in CZA compared to both PDB (one-sample t-test, p 2e-4) and MS (one-sample t-test, p 5e-8) (Fig. 3C). Similarly, genes present in shared H3K27me3 domains for CZA and PDB growth (111 genes) are not differentially expressed between CZA and PDB (one-sample t-test, p 0.92) when the genes are associated with H3K27me3, they are higher expressed in MS than in PDB (one-sample t-test, p 4e-3), but not higher expressed in MS than CZA (one-sample t-test, p 0.99). For genes present in shared H3K27me3 domains between MS and CZA growth (223 genes), we did not observe statistically significant transcriptional differences between the growth conditions where the genes lacked H3K27me3 domains (Fig. 4A, C). Analysing H3K27me3 domains unique to a medium, we found that CZA growth had the greatest number and proportion of unique genes locating in H3K27me3 domains (23.3%), followed by 8.0% unique to MS growth, and 1.7% unique to PDB growth (Fig. 3A). The genes uniquely marked in any condition did not show consistently increased expression in the condition in which the gene was not located in an H3K27me3 domain (Fig 3D). Overall these results suggest that differential expression can be associated with differential H3K27me3 domain status, but it is not a requirement. We observed clear examples where the loss of H3K27me3 in one medium is associated with increased transcription in that medium, but this was not universally true. Many genes undergo differential gene expression between growth conditions and remain in stable H3K27me3 domains. We note that the majority of genes located in H3K27me3 domains were common to all three growth conditions, accounting for 83.3%, 75.3% and 68.2% of the identified genes in H3K27me3 domains from PDB, MS and CZA growth, respectively, indicating that the qualitative presence of H3K27me3 domains is globally stable.

**FIG 3.**
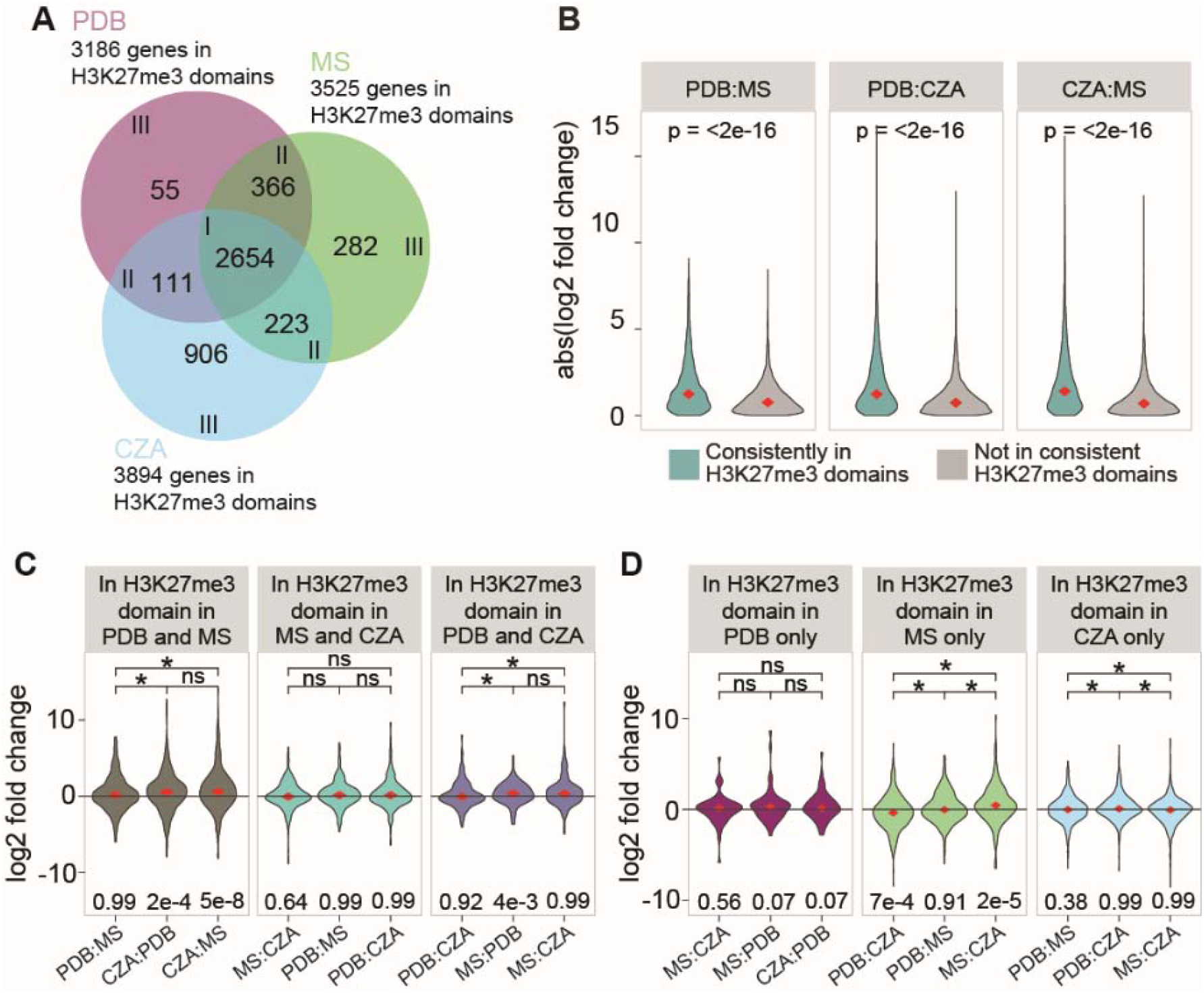
Differential H3K27 methylation partially explains differential expression. A) The number of genes located in H3K27me3 domains for *V. dahliae* cultivated for 6 days in Czapec-Dox (CZA), half-strength Murashige Skoog medium (MS) and potato dextrose broth (PDB). B) Absolute log2 fold-changes in expression between pairs of growth media. The genes were grouped between those consistently localized in H3K27me3 domains (2654 genes) and those not consistently present in H3K27me3 domains. C,D) Log2 fold-change values for each pair of growth medi1 for genes in H3K27me3 domains found in C) two or D) one of three tested *in vitro* growth media. Differential gene expression comparisons are set such that genes higher expressed in the medium in which they do not locate in H3K27me3 domain will have positive log2 fold-change. In case differential gene expression comparisons are between growth media in which the genes locate in H3K27me3 domains in C) both or D) neither of the compared media, the orientation of positive and negative log2 fold-change is arbitrary. Median values of violin plots are indicated with diamonds and shown above the plot. Significant difference of TPM values between growth media are determined with the One-Sample Wilcoxon Signed Rank Test (*: p <= 0.05). The one-sample two-sided T-test was used to test whether sample means significantly differ from 0, p-value shown below the plot.

**FIG 4.**
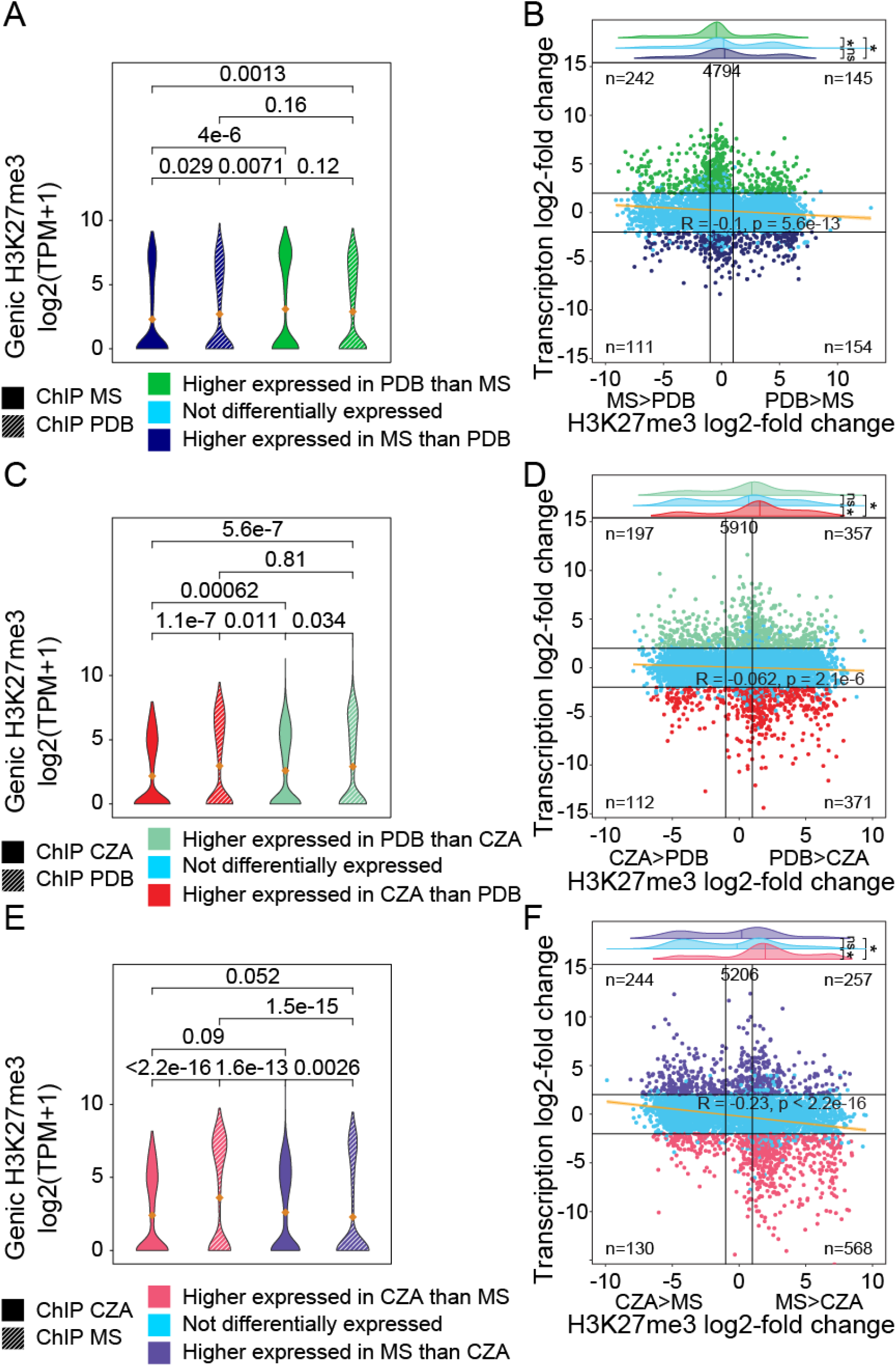
Differentially expressed genes associate with local changes in H3K27me3 coverage. Pairwise comparisons of input-corrected H3K27me3 coverage over differentially expressed genes for *V. dahliae* cultivated for 6 days in A,B) half-strength Murashige Skoog medium (MS) and potato dextrose broth (PDB), C,D) Czapec-Dox (CZA) and PDB, and E,F) MS and CZA. A,C,E) H3K27me3 coverage over differentially expressed genes in each corresponding growth medium. Mean values of violin plots are indicated with orange diamonds. B,D,F) per gene comparison of log_2_-fold change in transcription and genic H3K27me3 coverage, for genes with input-corrected H3K27me3 coverage >0 in either or both compared media (total number of genes indicated in top of plot). Black horizontal lines indicate transcription log_2_-fold change of −2 and 2, black vertical lines indicate H3K27me3 log_2_-fold change of −1 and 1. Numbers of genes in the four corner section are indicated. Significant difference of TPM values between growth media or gene sets are determined with the One-Sample Wilcoxon Signed Rank Test and indicated by their p-value or by asterisks (*: p≤0.05).

### Local quantitative differences in H3K27me3 levels are associated with transcriptional differences

Our analysis on H3K27me3 presence/absence dynamics did not account for potential quantitative differences between growth media. Whole chromosome plots of H3K27me3 domains identified between media reflect their generally stable presence (Fig. S8), but the analysis is based on qualitative H3K27me3 domain identification. Domain calling for H3K27me3 results in broad domains, but this fails to capture higher resolution quantitative differences that may exist between media, as is seen during inspection of global chromosome plots (Fig. S8). To understand how qualitative domain calling impacts the analysis, we examined quantitative differences between H3K27me3 ChIP-seq signal and transcriptional output between pairs of growth conditions. Genes were grouped based on differential gene expression between media, where genes that are significantly higher expressed in media A were one group and genes significantly that are higher expressed in media B another group. Subsequently, the H3K27me3 ChIP-seq signal relative to the input samples were normalized and compared between both growth media for the groups of differentially expressed genes (Fig. 4). Comparing results for PDB versus MS, we see that genes that are higher expressed in MS have a significantly lower MS H3K27me3 ChIP-signal versus the same genes from PDB ChIP (Fig. 4A). The contrasting comparison, genes that higher expressed in PDB, shows these genes do not significantly differ in ChIP signal between PDB and MS growth (Fig. 4A). Further integrating the transcriptional and ChIP fold-changes, for the 4,794 genes that have an input-corrected H3K27me3 ChIP signal above 0 in either MS and PDB, we see that genes higher expressed in PDB have a significantly lower log2 fold-change for H3K27me3 coverage, indicating these genes have lower H3K27me3-signal in PDB when compared with MS (Fig. 4B). Quantifying the number of genes in quadrant II and IV, we find that 396 (242+154) display a negative association between transcription and H3K27me3 ChIP-signal, whereas 256 (145+111) display a positive association (Fig. 4B). Thus, more genes are present in the quadrants that represent genes having lower H3K27me3 levels and higher transcription between the two growth conditions. The linear regression based on all genes is R=−0.1, also suggesting the low but significant negative association (Fig. 4B). It is clear that the association between differential expression and changes in ChIP-signal is not true for all genes, but overall we observe that the majority of genes that display changes for H3K27me3 ChIP-signal between media show the predicted transcriptional response where less H3K27me3 is associated with higher transcription (Fig. 4B).

Results for PDB versus CZA growth showed that genes higher expressed in CZA have higher H3K27me3 levels in PDB, consistent with the expected association (Fig. 4C). For genes higher expressed in PDB, however, we also observed a higher H3K27me3 level in PDB. Globally, the data indicated that genes higher expressed in CZA have more H3K27me3 ChIP signal in PDB than genes that are higher expressed in PDB, indicating a negative association between transcription and H3K27me3 presence (Fig. 4D). This is corroborated by the number of genes per quadrant, as 568 (197+371) genes locate in quadrants II and IV, whereas 469 (112+357) genes locate in quadrants I and III (Fig. 4D). The linear regression analysis indicates a slight negative association between differential expression and H3K27me3 ChIP-signal (Fig. 4D). We also observed an overall higher ChIP-signal from samples grown in CZA, but the reason for this is not clear.

Results for CZA versus MS growth showed that genes higher expressed in CZA had statistically higher levels of H3K27me3 ChIP-signal in MS medium, whereas genes higher expressed in MS have higher levels of H3K27me3 in CZA medium (Fig. 4E). The same pattern was observed in the integrated analysis, as genes higher expressed in CZA have more H3K27me3 signal in MS (Fig. 4F). There are 812 (244+568) genes in quadrant II and IV, supporting a negative association between differential transcription and H3K27me3 ChIP-signal, whereas 387 (130+257) genes are present in quadrants I and III. This is supported by the significant negative correlation (R=-0.23) (Fig. 4F). Overall, the results of the integrated analyses show that there is an association between quantitative transcriptional levels and H3K27me3 signal, where genes that are more highly expressed in a transcriptionally permissive medium have less H3K27me3 when compared with H3K27me3 levels in the repressive media. There are also many genes that were differentially expressed without an accompanying shift in H3K27me3 log2 fold-change (Fig. 4B,D,F). Collectively, these results are consistent with the observations of the qualitative H3K27me3 presence/absence comparisons, and suggest that H3K27me3 levels are generally stable, but local changes at particular genes may contribute to transcriptional dynamics.

### Genes induced *in planta* are largely H3K27me3-associated across all tested growth conditions

The presented analyses compared the directionality of H3K27me3 and transcriptional changes between axenic growth media to address if H3K27me3 dynamics are associated with transcriptional dynamics. Another interesting growth condition for *V. dahliae* is host colonization, and the contrast for the genes differentially expressed *in planta* when compared with axenic culture. We have thus far been unable to perform ChIP on *V. dahliae* during host colonization due to a low pathogen-to-plant biomass, and therefore cannot compare H3K27me3 levels between these conditions. However, we can identify the genes that are significantly induced during infection when compared with *in vitro* growth media and assess their chromatin profiles in these media to assess if *in planta*-induced genes appear heterochromatic during *in vitro* growth. Genes were grouped based on differential expression between PDB and *in planta* growth and chromatin profiles showed that genes that are higher expressed *in planta* had significantly more H3K27me3 during PDB growth when compared with genes that are more highly expressed in PDB (Fig. 5A). These *in planta*-induced genes lacked H3K4me2 and the DNA was less accessible during PDB growth (Fig. 5A). The association between *in planta* induction and higher H3K27me3 levels was not only seen relative to PDB growth, but also to growth in MS and CZA (Fig. 5B,C). For each media comparison, the genes that are more highly expressed *in planta* have higher H3K27me3 levels during axenic growth. To understand what genes are driving these differences, we investigated whether *in planta*-induced genes that locate in H3K27me3 domains are overrepresented for genes that are potentially involved in infection. There were approximately 600 genes that are more highly expressed *in planta* when compared with any of the three media (Fig. 5D). We found that, depending on the medium, from 41.2% to 52.8% of the *in planta*-induced genes were located in H3K27me3 domains (Fig. 5D). We observed that the *in planta*-induced genes within H3K27me3 domains have a higher, yet not statistically significant, fraction of genes encoding secreted proteins or putative effectors when compared with all *in planta*-induced genes. For example, genes in H3K27me3 domains that were *in planta*-induced when compared with PDB growth (277 genes total), 15.5% had a secretion signal and 4.3% were predicted effectors. This is compared to all *in planta*-induced genes relative to PDB (673 genes total), where 12.3% had a secretion signal and 2.7% were predicted effectors. These results were similar between the other two media, and we conclude that *in planta*-induced genes in H3K27me3 domains have a slightly higher fraction of genes that are potentially involved in infection when compared with *in planta*-induced genes not in H3K27me3 domains. Collectively, these results indicate that genes that are higher expressed *in planta* have a heterochromatic profile (i.e. H3K27me3 association, low H3K4me2, lower accessibility) when analysed in axenic culture. The chromatin profile of these genes during *in planta* growth will need to be directly assessed in the future.

**FIG 5.**
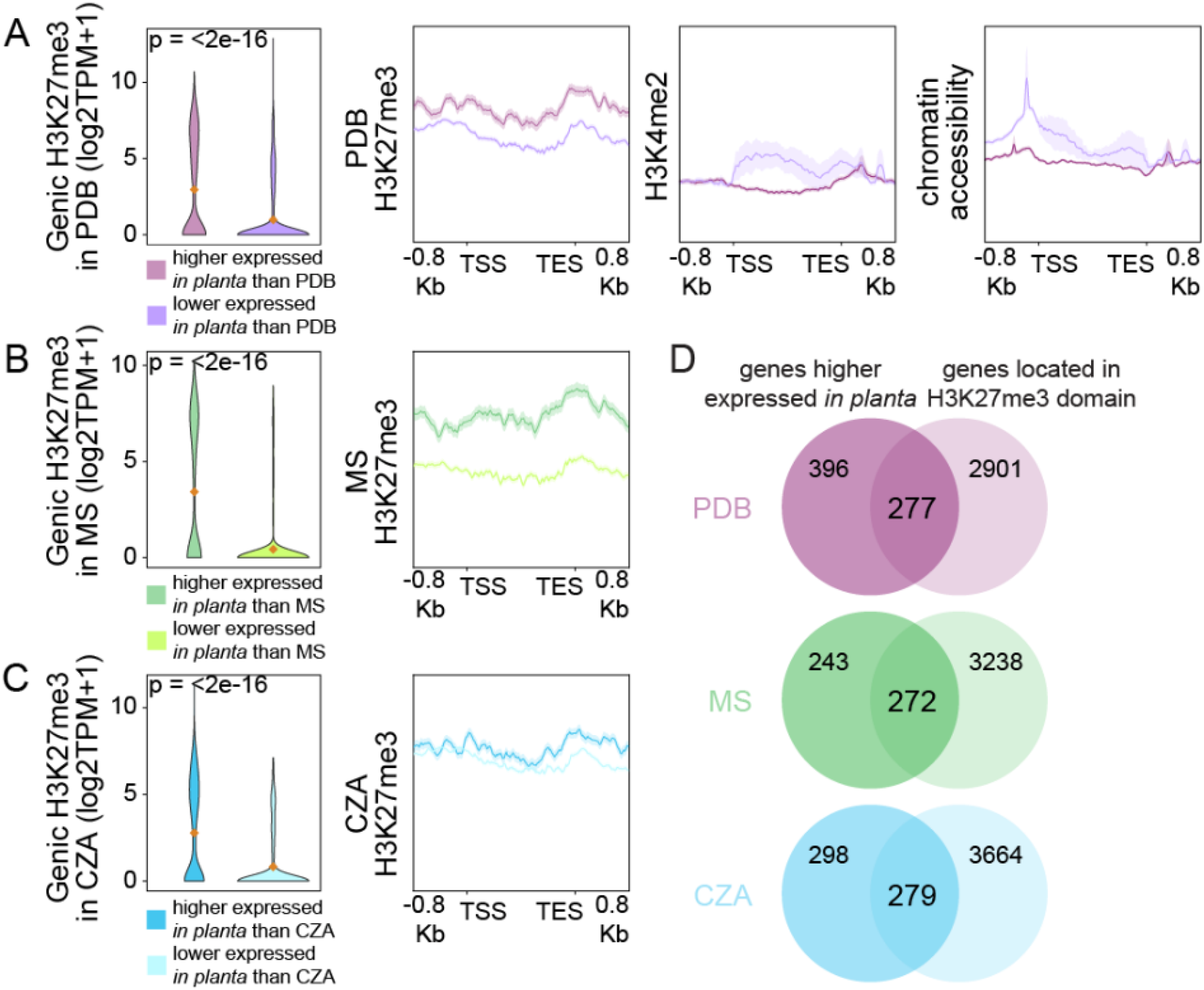
*In planta*-induced genes are H3K27me3-associated under transcriptionally repressive conditions. Violin plots display input-corrected H3K27me3 ChIP signal over genes differentially expressed between *in planta* and A) 6 days of cultivation in potato dextrose broth (PDB), B) half strength Murashige Skoog medium (MS), and C) Czapec-Dox medium (CZA). Mean values are indicated by orange diamonds. Line plots display average coverage of chromatin features over the gene sets. D) overlap of genes that are higher expressed *in planta* than in PDB, MS or CZA with genes located in an H3K27me3 domain in the corresponding growth medium. Significant differences of H3K27me3 coverage are determined with the One-Sample Wilcoxon Signed Rank Test (*: p <= 0.05).

## DISCUSSION

Chemical histone modifications play an essential role in transcriptional regulation, but a number of mechanistic questions remain for their role in differential transcription in filamentous fungi. In this study, we show that many genes that are differentially transcribed between *in vitro* growth conditions are located in H3K27me3 domains. This is interesting because the mark contributes to transcriptional repression and it is relatively sparse in the genome, occupying approximately 33% of the genes. However, 35% to 70% of differentially expressed genes identified between growth conditions reside within H3K27me3 domains. It is not clear what mechanism might drive this association.

The global fraction of genes that locates in H3K27me3 domains in *V. dahliae* is similar to what has been reported for the asocomycete *Podospora anserina* (28), but higher than for *N crassa, Zymoseptoria tritici, Leptosphaeria maculans*, and *Magnaporthe oryzae* (9-16% of genes in H3K27me3 domain) (20, 30, 32, 50). Previous reports showed that H3K27me3 represses transcription of secondary metabolite clusters in *Fusarium spp.* and in *Epichloe festucae*, and of effectors in *Z. tritici* and *M. oryzae* (22, 24, 31, 32, 50). However, these reports have mainly relied on genomic association with H3K27me3, genetic perturbation altering H3K27me3 deposition, and the analysis of only few genes under natural conditions. Given our results, and these previous findings, our hypothesis was that differences in transcription between *in vitro* growth conditions is in part coordinated by dynamics for H3K27 methylation. We directly addressed this phenomenon using genome-wide H3K27me3 profiling and RNA-seq under different *in vitro* growth conditions. Consistent with reports in other fungi, we see that H3K27me3 is associated with transcriptional repression, and that deleting *Set7*, encoding the histone methyltransferase that is responsible for H3K27me3, led to induction of many genes normally present in H3K27me3 domains. Importantly, our direct observations of H3K27me3 levels and transcriptional output from three *in vitro* media showed that H3K27me3 domains are generally stable globally. We see that 50% to 75% of the identified H3K27me3 domains, depending on the medium analyzed, did not change between the tested conditions. Despite many H3K27me3 domains not undergoing presence/absence dynamics, numerous genes in these domains displayed differential transcription between the tested conditions, indicating that complete loss of H3K27me3 is not strictly required for transcriptional induction.

One possibility to account for these seemingly contradictory results is that even though H3K27me3 appears stable at the level of broad-peak calling, there may be smaller regions within the broad domain that are dynamic. Assessing quantitative differences in H3K27me3 ChIP-seq levels at defined genomic locations between paired *in vitro* growth conditions indicates that this can account for some of the observed transcriptional differences. For example, some genes have a lower H3K27me3 ChIP-signal upon growth in transcriptionally permissive medium when compared with growth in a transcriptionally repressive medium. We interpret these results as evidence for local, rather than global, changes in H3K27me3 dynamics contributing to transcriptional regulation for the underlying genes. H3K27me3-associated genes that are differentially expressed while presence of the histone mark remains stable, may be transcriptionally regulated through the activity of H3K27me3 readers. For instance, the *Fusarium graminearum* histone reader BP1, which is orthologous to *N. crassa* EPR-1, specifically binds to H3K27me3 and co-represses gene transcription (51, 52). The gene encoding BP1 is conserved within fungi, including *V. dahliae* (51). Dynamic binding of such transcription-repressing histone reader to stable H3K27me3 domains may explain the observed transcriptional dynamics.

We note that an individual histone does not permit for a quantitative tri-methylation status, as an individual H3K27 is either tri-methylated or not. At the nucleosome level, there can be none, one, or two H3K27 tails with a tri-methyl modification. At the cell population level, there can be considerable quantitative differences because of heterogeneity for histone modification status between individual cells. Cell variability may be the source of quantitative differences observed in our experiments, arising from variation for both the percent of cells with H3K27me3 and the number of tails of a nucleosome with H3K27me3. While we cannot determine this based on our present data, our results show that some genes that are differentially expressed between growth conditions are associated with quantitative differences in H3K27me3 levels, providing evidence that chemical histone modifications dynamics can be involved in a transcriptional regulatory mechanism in *V. dahliae*. The concept of local versus global H3K27me3 changes is consistent with data from *M. oryzae*, in which direct *in planta* H3K27me3 ChIP-qPCR showed that some, but not all, analyzed regions displayed differential H3K27me3 levels consistent with increased transcription between *in planta* and *in vitro* conditions (50). We have not been able to directly assess histone modifications for *V. dahliae* during plant infection, but we did analyze the status of H3K27 methylation during *in vitro* growth of *in planta*-induced genes. The *in planta*-induced genes have significantly higher levels of H3K27me3 and the DNA is substantially less accessible during *in vitro* growth when compared with genes that are highly expressed during *in vitro* growth. We conclude that dynamics for H3K27me3 can contribute to differential gene expression under natural conditions, but our results show this is not required. Our conclusions are limited to the tested conditions, and it is possible that analyzing H3K27me3 levels in different cell types or growth stages (e.g. spores, microsclerotia, *in planta* infection) may yield a different picture of global H3K27me3 distribution. Additionally, our experiments focused on steady-state growth on different media as these provide a clear and reproducible transcriptional difference. It is possible that H3K27me3 dynamics occur rapidly in response to environmental changes, cues, or developmental stages that we did not capture.

It appears evident that H3K27me3 contributes to additional genome functions beyond strict transcriptional repression, at least in fungi (37). This is supported by our data showing that the majority of H3K27me3 domains were stable between the tested growth conditions, indicating these H3K27me3 domains have additional functions. Stable H3K27me3 domains may represent another form of constitutive heterochromatin, but this fact seems unlikely given that in our data many stable H3K27me3 domains harbored differentially expressed genes between growth media. Additionally, results in *N. crassa* have shown that genetic perturbation leading to the genome-wide loss of H3K9me3 causes a redistribution of H3K27me3 to previously H3K9me3 marked sites, but interestingly, this re-distribution does not result in transcriptional silencing (53). Rather, the redistribution of H3K27me3 in *N. crassa* lacking H3K9me3 appears to contribute to genomic instability (53, 54). In *Z. tritici*, the enrichment of H3K27me3 at dispensable chromosomes and empirical evidence that H3K27me3 somehow increases genomic instability, additionally supports the hypothesis that H3K27me3 contributes to additional genomic functions beyond transcriptional regulation (32, 48). Evolutionary analysis across *Fusarium* and related species indicates that genes marked by H3K27me3 have a higher duplication rate in *Fusarium* and are less conserved with more distantly related species (55). Additional research is needed to fully understand the mechanisms of H3K27me3 targeting, dynamics, and impact on genome stability in fungi. The results presented here show that H3K27me3 domains are largely similar between *in vitro* growth conditions, but that quantitative differences in H3K27me3 levels can be associated with concomitant transcriptional differences between the tested conditions.

## MATERIALS AND METHODS

### Fungal growth conditions

*Verticillium dahliae* strain JR2 (CBS 143773; Faino et al., 2015) was cultured on potato dextrose agar (PDA) (Oxoid, Thermo Scientific) at 22°C in the dark. Liquid cultures were obtained by collecting conidiospores from PDA plates after approximately 2 weeks of growth followed by inoculation at a final concentration of 1×10^4^ spores per mL into liquid growth media. Media used in this study are potato dextrose broth (PDB) (Difco, Becton Dickinson, Franklin Lakes, NJ, USA), half strength Murashige & Skoog plus vitamins (MS) (Duchefa-Biochemie, Haarlem, The Netherlands) medium supplemented with 3% sucrose, and Czapec-Dox medium (CZA) (Oxoid, Thermo Scientific, Waltham, MA, USA). Liquid cultures were grown for 6 days in the dark at 22°C at 140 RPM. Mycelium was collected by straining cultures through miracloth (22 μm) (EMD Millipore, Darmstadt, Germany) and pressing to remove liquid after which the mycelium was flash frozen in liquid nitrogen and ground to powder with a mortar and pestle. If required, samples were stored at −20°C prior to nucleic acid extraction. All analyses were performed based on triplicate cultures that were processed individually.

### RNA sequencing and analysis

RNA was isolated from ground mycelium using TRIzol (Thermo Fisher Science, Waltham, MA, USA) following manufacturer guidelines. Contaminating DNA was removed using the TURBO DNA-free kit (Ambion, Thermo Fisher Science, Waltham, MA, USA) and RNA integrity was assessed with gel-electrophoresis and quantified using a Nanodrop (Thermo Fisher Science, Waltham, MA, USA). Library were prepared and singleend 50 bp sequenced on the DNBseq platform at BGI (Hong Kong, China).

Sequencing reads were mapped to the reference annotation of *V. dahliae* strain JR2 (56) using Kallisto quant (57) to calculate per gene TPM values. Differential expression between cultivation in PDB and CZA, MS or during colonization of *Arabidopsis thaliana* was determined using DESeq2 (58).

### ChIP-sequencing and analysis

ChIP-seq was performed as described previously (Cook et al., 2020). Frozen ground mycelium was added to ChIP lysis buffer, dounced 40 times and subsequently sonicated for 5 rounds of 20 seconds with 40 second rest stages on ice. Supernatants were collected after centrifugation and treated with MNase (New England Biolabs, Ipswich, MA, United States) for 10 minutes in a 37°C waterbath. MNase activity was quenched by addition of EGTA and samples were subsequently pre-cleared by addition of 40 µl Protein A Magnetic Beads (New England Biolabs, Ipswich, MA, United States) and rotating at 4°C for 60 min. Beads were captured and supernatant was divided over new tubes containing antibodies against either H3K4me2, H3K9me3 or H3K27me3 (ActiveMotif; #39913, #39765 and #39155) and incubated overnight with continuous rotation at 4°C. Subsequently, the antibodies were captured, washed and nucleosomes were eluted from beads, after which DNA was treated with Proteinase-K and cleaned-up using chloroform. DNA was isolated by overnight precipitation in ethanol and DNA concentration was determined with the Qubit™ dsDNA HS Assay Kit (Thermo Fisher Science, Waltham, MA, USA). Sequencing libraries were generated using the TruSeq ChIP Library Preparation Kit (Illumina, San Diego, CA, United States) according to instructions, but without gel purification and with use of the Velocity DNA Polymerase (BioLine, Luckenwalde, Germany) for 25 cycles of amplification. Single-end 125bp sequencing was performed on the Illumina HiSeq2500 platform at KeyGene N.V. (Wageningen, the Netherlands).

Raw reads were trimmed using TrimGalore (59) and mapped to the *V. dahliae* strain JR2 reference genome (56) using BWA-mem with default settings (60). Three regions of the genome were masked due to aberrant mapping, (chr1:1-45000, chr2:3466000-3475000, chr3:1-4200). ATAC-seq reads of *V. dahliae* cultured in PDB (44) were treated similarly as ChIP-seq reads, but the paired end read pairs were trimmed and mapped simultaneously, and only read-pairs <100 bp were considered for further analyses, as these represent open DNA. Mapped reads were RPGC-normalized using deepTools bamCoverage (61) with binsize 1,000 and smoothlength 3,000 for plotting over the genome, and binsize 10 and smoothlength 30 for further analysis.

Normalized replicate samples with high correlation were selected and mean datasets per growth media were generated with input controls for background signal correction using WiggleTools mean (62). H3K27me3-enriched regions were determined on selected replicates with input control using epic2 with a binsize of 2,500 bp (63). Average coverage over gene bodies per expression quintile was calculated using deepTools computeMatrix in scale-regions mode (61). Expression quintiles were generated by sorting all genes based on their average TPM value from three replicates of *V. dahliae* cultured for 6 days in PDB. We used deepTools multiBigwigSummary (61) to determine the presence of H3K27me3 over gene bodies (region between TSS and TES) for each replicate ChIP sample, as well as for input controls. Samples and input-controls were TPM normalized, after which the normalized input-control signal was subtracted from normalized H3K27me3 TPM values and resulting negative values were set to 0. Changes in H3K27me3 levels between growth media were determined by taking the average input-normalized H3K27me3 TPM values +1 and calculating the log2-fold change for each pair-wise comparison.

### Generation of *Set7* deletion mutant

The *Set7* deletion mutant (Δ*Set7*) was constructed as previously described (64). Briefly, genomic DNA regions flanking the 5’ and 3’ ends of the coding sequences were amplified with PCR using primers listed in Table S1 (primers 1-4) and cloned in to the pRF-HU2 vector (65) using USER enzyme following the manufacturer’s protocol (New England Biolabs, MA, USA). Sequence-verified vectors were transformed into *Agrobacterium tumefaciens* strain AGL1 and used for *V. dahliae* conidiospore transformation as described previously (64). *V. dahliae* transformants that appeared on hygromycin B were transferred to fresh PDA supplemented with hygromycin B after five days. Putative transformants were screened using PCR to verify deletion of the target gene sequence and integration of the selection marker at the designated locus using primers listed in Table S1 (primers 5-8). Analysis and comparison of gene expression in the *Set7* deletion mutant to type *V. dahliae* was performed as described above.

## ACKNOWLEDGEMENTS

This work was supported by a PhD grant of the Research Council Earth and Life Sciences (project 831.15.002) to HMK, and by Human Frontier Science Program Postdoctoral Fellowship (HFSP, LT000627/2014-L), by USDA’s National Institute of Food and Agriculture (award no. 2018-67013-28492) through the Plant Biotic Interactions Program, and by the National Science Foundation (award no. 1936800) Division of Molecular and Cellular Biosciences to DEC. Work in the laboratories of M.F.S and B.P.H.J.T. is supported by the Research Council Earth and Life Sciences (ALW) of the Netherlands Organization of Scientific Research (NWO). B.P.H.J.T acknowledges funding by the Alexander von Humboldt Foundation in the framework of an Alexander von Humboldt Professorship endowed by the German Federal Ministry of Education and Research is furthermore supported by the Deutsche Forschungsgemeinschaft (DFG, German Research Foundation) under Germany’s Excellence Strategy – EXC 2048/1 – Project ID: 390686111.

## ADDITIONAL FILES

### Supplementary Figures

**Fig. S1.** Distribution of chromatin features over all chromosomes.

**Fig. S2.** Gene expression correlates with histone modification presence and chromatin accessibility.

**Fig. S3.** Western blot shows loss of H3K27me3 in the *V. dahliae* Δ*Set7* deletion mutant.

**Fig. S4.** ChIP-sequencing shows loss of H3K27me3 in the *V. dahliae* Δ*Set7* mutant.

**Fig. S5.** Genes associated with H3K27me3 in wild type *V. dahliae* are stronger transcriptionally induced in the Δ*Set7* mutant than not H3K27me3 associated genes..

**Fig. S6.** Correlation between H3K27me3 ChIP samples

**Fig. S7.** Differentially expressed genes are enriched in H3K27me3 domains during cultivation in the non-permissive growth medium.

**Fig. S8.** Distribution of H3K27me3 for *V. dahliae* cultivated in *in vitro* growth media.

### Supplementary Tables

**Table S1.** Primers used to delete and analyse the *Set7* coding sequence in *V. dahliae*.

