## Supplemental Data for "Local rather than global H3K27me3 dynamics associates with differential gene expression in *Verticillium dahliae*"

### Supplementary Data

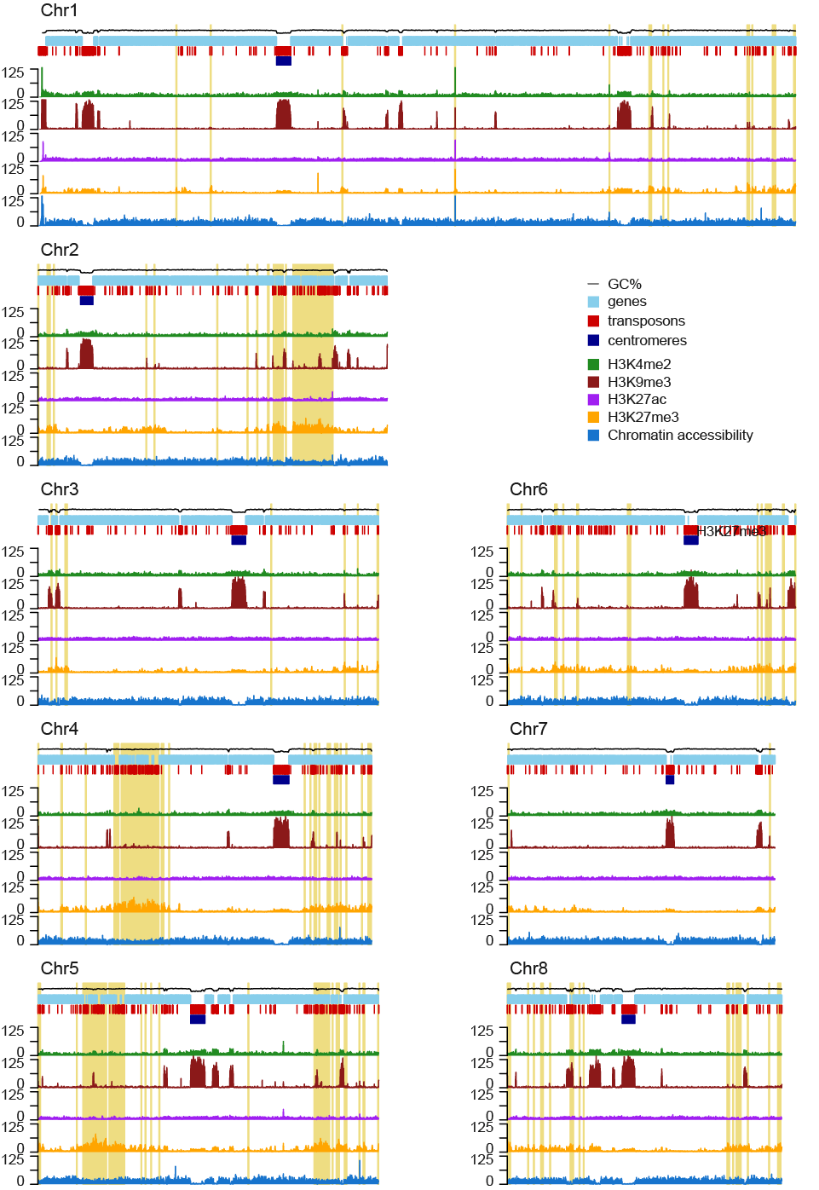

**Figure S1. Distribution of chromatin features over all chromosomes.** Whole genome distribution of the euchromatin-associated histone modification H3K4me2 (green line), the constitutive heterochromatin-associated histone modification H3K9me3 (red line), the facultative heterochromatin-associated histone modification H3K27me3 (yellow line) and chromatin accessibility (blue line) as determined by ATAC-seq. Genes are indicated in light blue, transposons are indicated in red, centromere is indicated in dark blue and adaptive genomic regions are indicated in yellow.

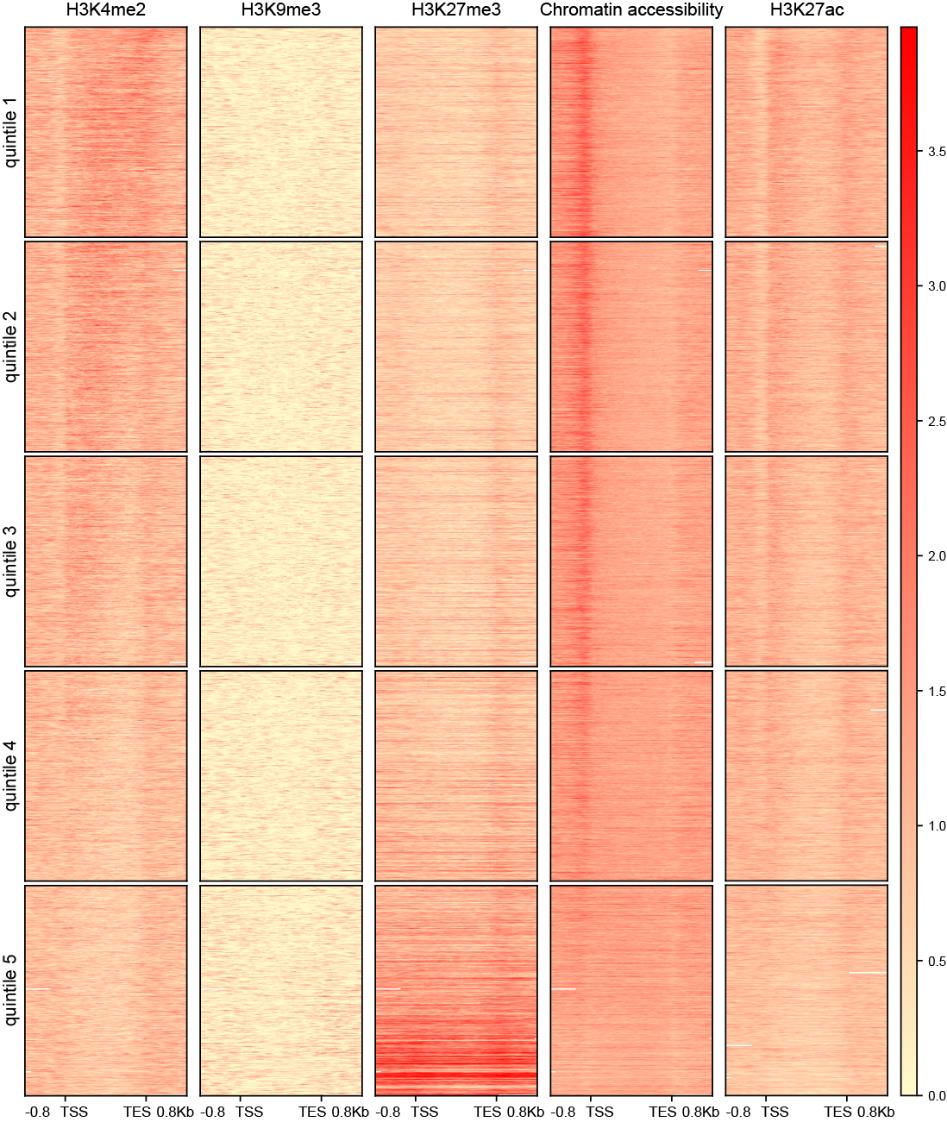

**Figure S2. Gene expression correlates with histone modification presence and chromatin accessibility.** RPGC normalized coverage of the histone marks H3K4me2, H3K9me3, H3K27me3 and chromatin accessibility over gene bodies (between transcription start site (TSS) and transcription end site (TES)) ±800 bp of flanking sequence. Each row represents a single gene, and are sorted based on their TPM value upon cultivation for six days potato dextrose broth (PDB), with the top gene being most highly expressed. Genes are grouped in expression quintiles as in Figure 3A.

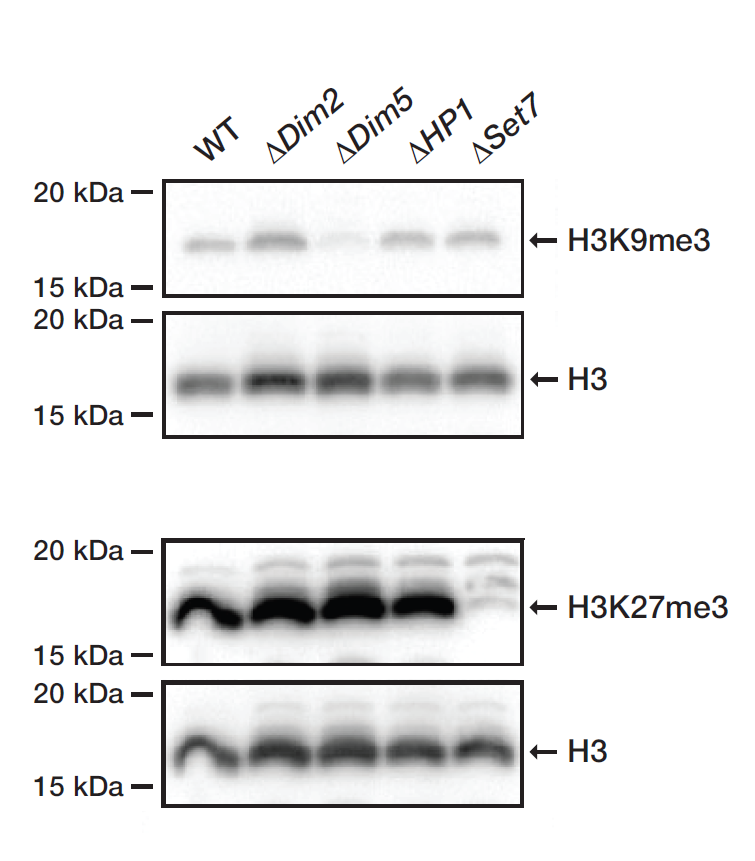

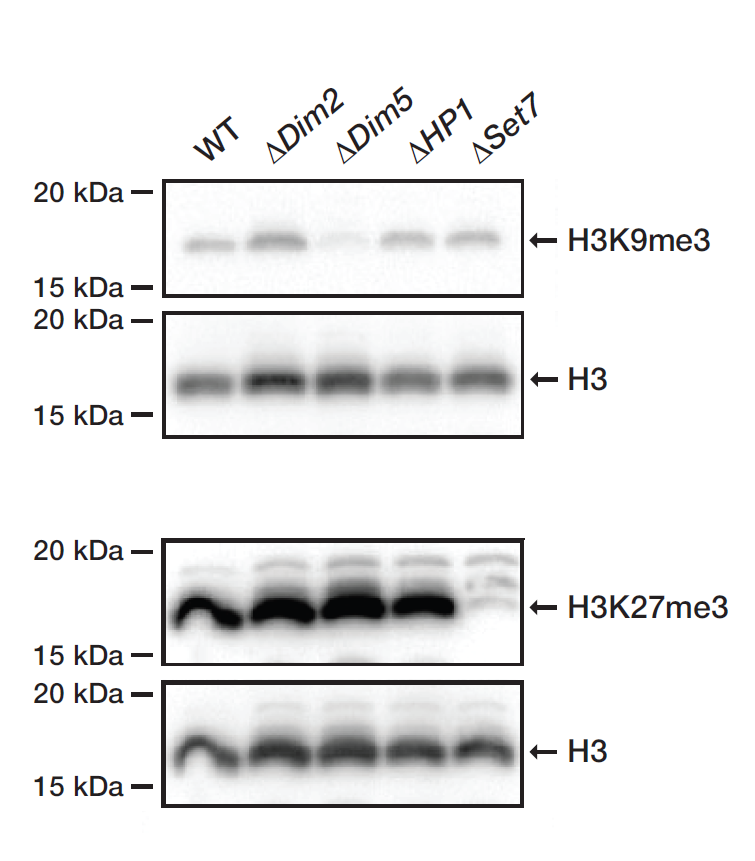

**Figure S3. Western blot shows loss of H3K27me3 in the V. dahliae Set7 deletion mutant.** Histone isolations of wild-type, ∆Dim2, ∆Dim5, ∆HP1 and ∆Set7 were tested for presence of the H3K27me3 histone modification by Western blotting. The antibody against H3 was used as loading control.

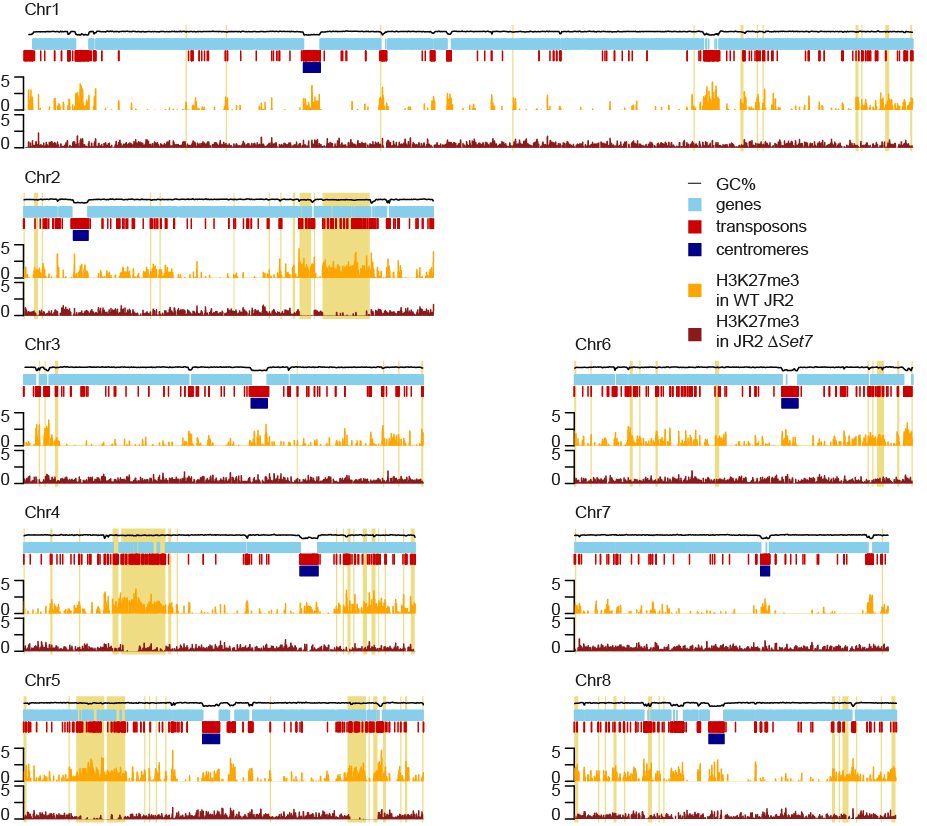

**Figure S4.** **ChIP-sequencing shows loss of H3K27me3 in the V. dahliae ∆Set7** **mutant.** H3K27me3 ChIP coverage over the genome in a triplicate of JR2 WT (yellow) and in a duplicate of JR2 ∆Set7, cultivated for 6 days in potato dextrose broth.

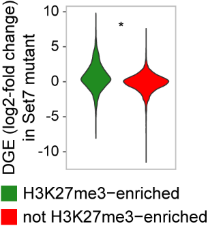

**Figure S5.** **Genes associated with H3K27me3 in wild type V. dahliae are stronger transcriptionally induced in the ∆Set7** **mutant than not H3K27me3 associated genes.** Log2-fold change of expression between wild type and ∆Set7 mutant for genes associated with H3K27me3 in wild type (green) and those not associated with H3K27me3 in wild type (red). Significant difference of log2-fold change between gene sets are determined with the One-Sample Wilcoxon Signed Rank Test (*: p <= 0.05).

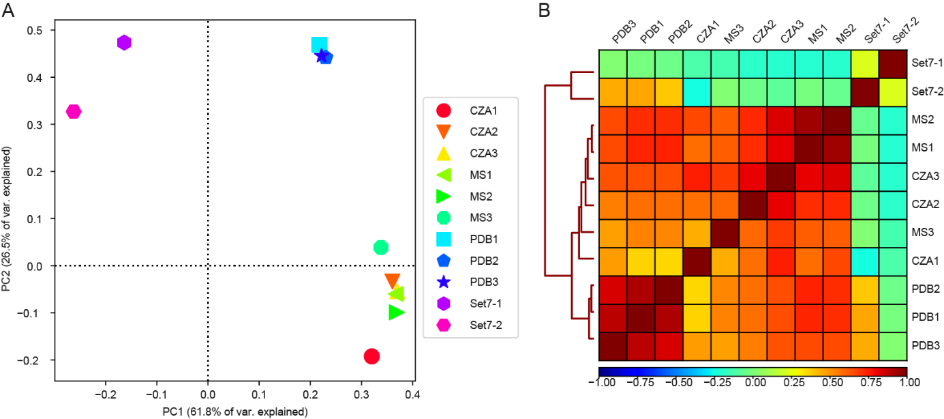

**Figure S6.** **Correlation between H3K27me3 ChIP samples.** A) PCA plot and B) heatmap displaying between-sample correlation of H3K27me3 coverage for 1kb bins for triplicates of JR2 WT cultivated for 6 days in Czapec-Dox medium (CZA), half-strength Murashige-Skoog medium (MS) and potato dextrose broth (PDB), and a duplicate JR2 ∆Set7 cultivated for 6 days in PDB.

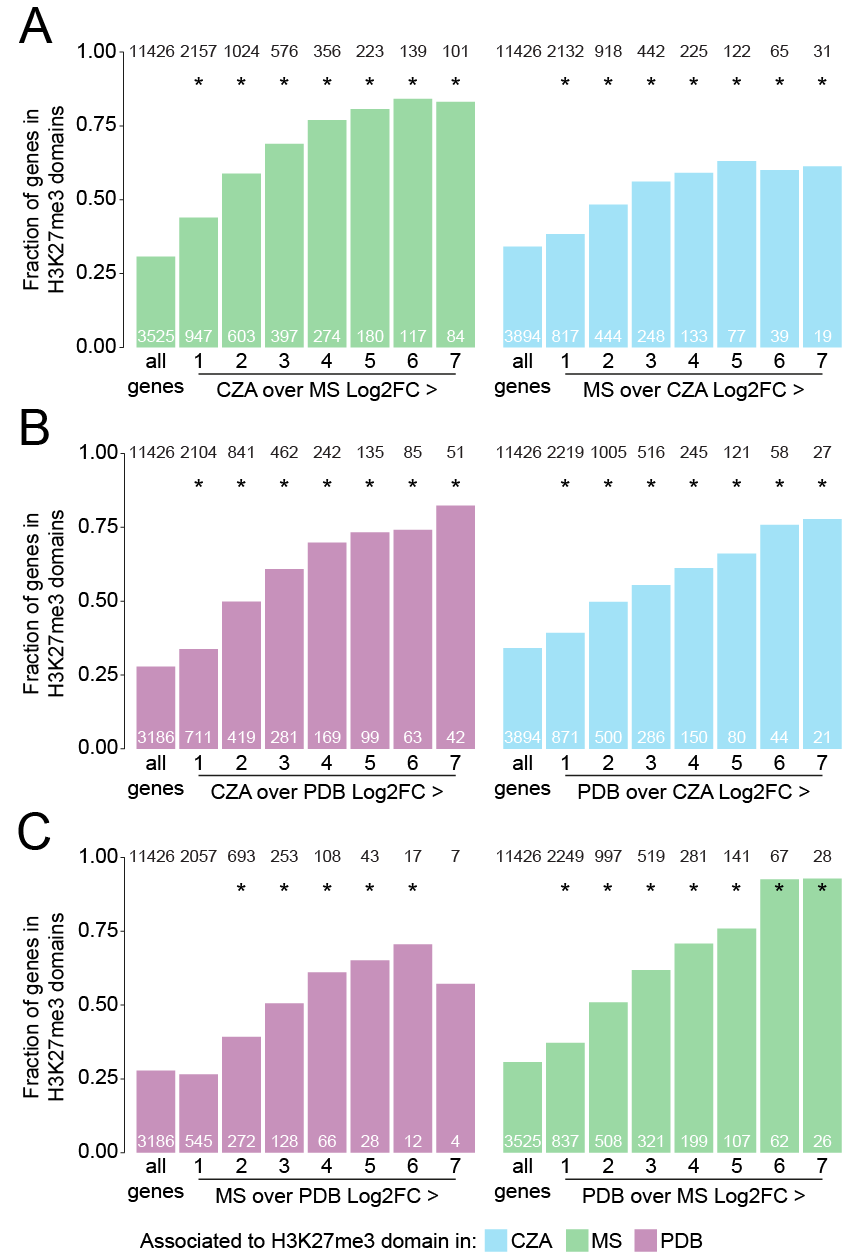

**Figure S7****. Differentially expressed genes are enriched in H3K27me3 domains during cultivation in the non-permissive growth medium.** Fractions of total and differentially expressed genes (Log2-fold changes >1, 2, 3, 4, 5, 6 and 7) that are associated with an H3K27me3 domain in the non-permissive condition Czapec-Dox medium (CZA, blue bars), half-strength Murashige-Skoog medium (MS, green bars) or Potato-Dextrose broth (PDB, red bars) between (A) cultivation for 6 days in CZA and MS, (B) cultivation for 6 days in CZA and PDB and (C) cultivation for 6 days in MS and PDB. Black numbers above the bars indicate numbers of genes per gene set. White numbers at bottom of the bars indicate the numbers of genes that locate in an H3K27me3-domain per gene set. Significance of fraction differences was calculated for each fraction of higher expressed genes in adaptive genomic regions when compared with the fraction of total genes in adaptive genomic regions, by the two-sided two-proportions Z-test (*: p$\leq$0.05).

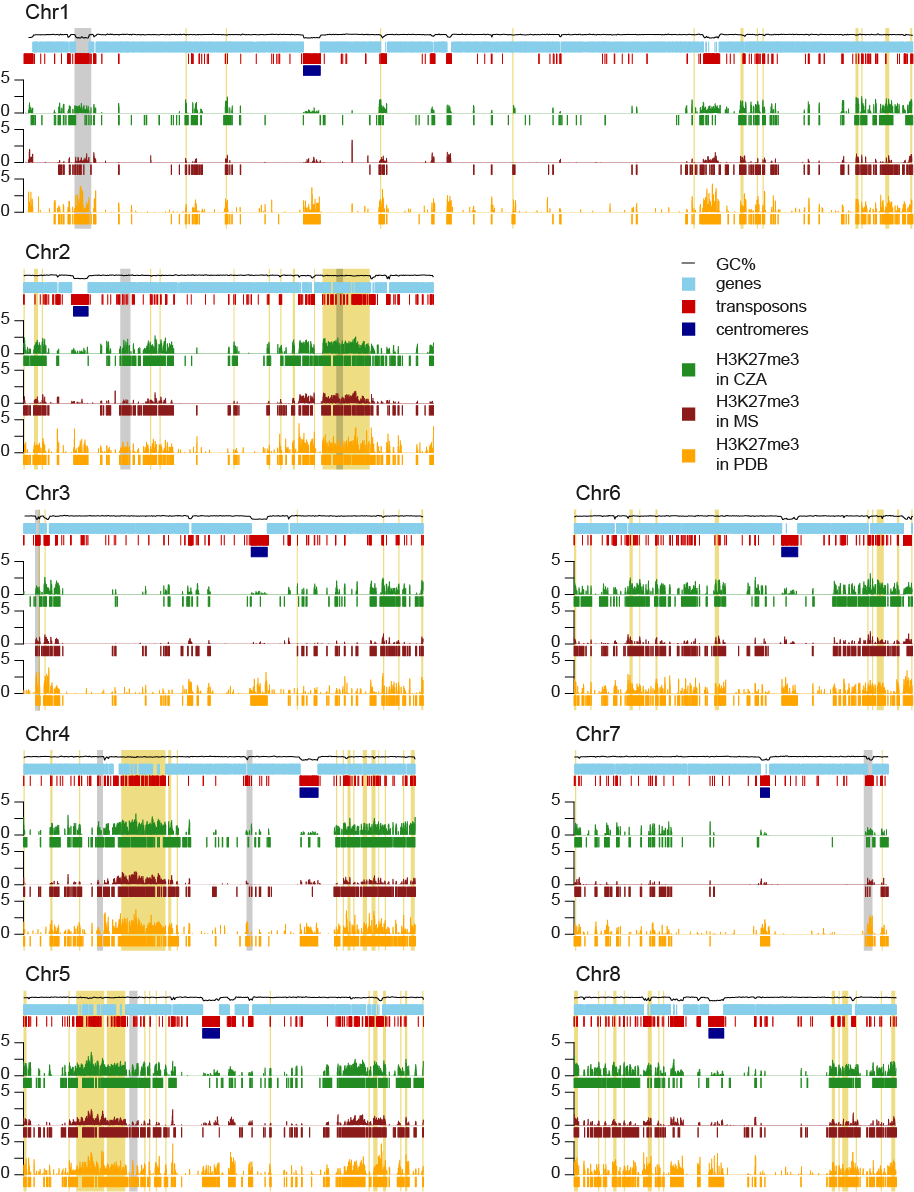

**Figure S8.** **Distribution of H3K27me3 for V. dahliae cultivated in vitro.** Average H3K27me3 distribution for replicates of V. dahliae cultivated for 6 days in potato dextrose broth (PDB, indicated in yellow), half strength Murashige Skoog medium (MS, indicated in red) and Czapec-Dox medium (CZA, indicated in green). Predicted H3K27me3 domains are indicated as blocks below each H3K27me3 track. Adaptive genomic regions are highlighted in yellow. Genomic regions within H3K27me3 domains that are visibly different between growth conditions are highlighted in grey.

**Table S1: Primers used to delete and analyze the Set7 coding sequence in V. dahliae.**

| **Name** | **Sequence** | **Purpose** | **Number** |
| --- | --- | --- | --- |
| Set7.ko-LB_F | *GGTCTTAAU*TGAGCTTGACAGTTCAGTTGTCG | Amplify left flanking sequence | 1 |
| Set7.ko-LB_R | *GGCATTAAU*AAGTTGTGTTGTCAGCGTGCATA | Amplify left flanking sequence | 2 |
| Set7.ko-Rb_F | *GGACTTAAU*ATCAAGTCCGCCTACTTTCCAAG | Amplify right flanking sequence | 3 |
| Set7.ko-Rb_F | *GGGTTTAAU*GTGGAGAATCGTCTGGGGTTATC | Amplify right flanking sequence | 4 |
| Set7_Confirm_F | CCTCCAGCTCCTGAAGAAGAA | Confirm gene replacement (selection) | 5 |
| Vector_Reverse | GGAGTCGCATAAGGGAGAGCG | Confirm gene replacement (selection) | 6 |
| Set7.ORF.270_F | TCACAGCCGCTACATCAATCA | Confirm ORF is absent in KO | 7 |
| Set7.ORF.270_R | TCGTGTTGAACCTCCTTGGAC | Confirm ORF is absent in KO | 8 |
| Italic sequences at the 5' ends represent adapters added for USER cloning | | | |
